# *Legionella pneumophila* targets autophagosomes and promotes host autophagy during late infection

**DOI:** 10.1101/2021.07.08.451723

**Authors:** Rebecca R. Noll, Colleen M. Pike, Stephanie S. Lehman, Chad D. Williamson, M. Ramona Neunuebel

**Author notes:** Correspondence to M. Ramona Neunuebel ORCID: 0000-0002-3113-7144. C.M.P and R.R.N. contributed equally to this work.

## Abstract

Autophagy is a fundamental eukaryotic process that mediates clearance of unwanted molecules and facilitates nutrient release. The bacterial pathogen *Legionella pneumophila* establishes an intracellular niche within phagocytes by manipulating host cellular processes, such as autophagy. Effector proteins translocated by *L. pneumophila*’s Dot/Icm type IV secretion system have been shown to suppress autophagy. However evidence suggests that overall inhibition of autophagy may be detrimental to the bacterium. As autophagy contributes to cellular homeostasis and nutrient acquisition, *L. pneumophila* may translocate effectors that promote autophagy for these benefits. Here, we show that effector protein Lpg2411 binds phosphatidylinositol-3-phosphate lipids and preferentially binds autophagosomes. Translocated Lpg2411 accumulates late during infection and co-localizes with the autophagy receptor p62 and ubiquitin. Furthermore, autophagy is inhibited to a greater extent in host cells infected with a mutant strain lacking Lpg2411 compared to those infected with wild-type *L. pneumophila,* indicating that Lpg2411 stimulates autophagy to support the bacterium’s intracellular lifestyle.

**Summary:** *Legionella pneumophila* translocates several effector proteins that inhibit autophagic processes. In this study, we find that the effector protein Lpg2411 targets autophagosomes during late stages of infection and promotes autophagy.

## Introduction

To survive within eukaryotic cells intracellular bacterial pathogens must be able to acquire nutrients and evade degradation by processes like autophagy. In eukaryotes, autophagy is a well-conserved pathway involved in clearance of long-lived or damaged proteins and the recovery of nutrients. Bacterial pathogens can be engulfed by autophagosomes and targeted for lysosomal degradation, an autophagic process known as xenophagy (Kohler and Roy, 2017). Many pathogens that proliferate within eukaryotic cells, including *Legionella pneumophila*, have acquired the ability to manipulate autophagic processes for their survival (reviewed in (Siqueira et al., 2018)).

*Legionella pneumophila* is an intracellular opportunistic pathogen and an important cause of hospital- and community-acquired pneumonia (Mudali et al., 2020). To infect and replicate within phagocytic cells, *L. pneumophila* has evolved a complex survival strategy that relies on the establishment of a protective membrane-bound vacuole that avoids the endolysosomal pathway and host surveillance system (Roy and Tilney, 2002). Once engulfed into a phagosome, *L. pneumophila* directs membrane remodeling of the phagosomal compartment into an endoplasmic reticulum (ER)-like compartment, termed the *Legionella*-containing vacuole (LCV), by recruiting ER-to-Golgi transport vesicles (Hubber and Roy, 2010; Kagan and Roy, 2002; Swanson and Isberg, 1995).

LCV formation and maintenance depends on a type IV secretion system (T4SS) named Dot/Icm, which translocates over 350 effector proteins into the host cell (Roy et al., 1998; Steiner et al., 2018; Vogel et al., 1998). These effectors modulate prominent cellular pathways, such as endosomal maturation, ubiquitination and autophagy to promote intracellular survival (Qiu and Luo, 2017a; Qiu and Luo, 2017b; Thomas et al., 2020). *L. pneumophila* inhibits host autophagy by translocating multiple effector proteins that interfere with the formation and maturation of autophagosomes (Choy et al., 2012; Rolando et al., 2016; Thomas et al., 2020). However, *L. pneumophila* replication is hindered when autophagosome formation is impaired (Amer and Swanson, 2005; Sturgill-Koszycki and Swanson, 2000), suggesting that autophagy may confer some benefit for the intracellular survival of *L. pneumophila* and that additional effectors counterbalance this degradative flux pathway.

Autophagic clearance of proteins destined for lysosomal degradation requires the actions of autophagic receptors (Johansen and Lamark, 2011). One such receptor, p62/SQSTM1, functions as an adaptor between ubiquitinated proteins and LC3 on the autophagosome membrane (Kirkin et al., 2009; Komatsu et al., 2007). p62 is an essential component for xenophagic targeting and degradation of important bacterial pathogens such as *Salmonella enterica* and *Listeria monocytogenes* (Yoshikawa et al., 2009; Zheng et al., 2009). The investigation of p62 during *L. pneumophila* infection has yielded conflicting results. p62 was shown to target the LCV for autophagosomal degradation in restrictive bone marrow-derived macrophages at 1, 2, and 6 hours post-infection (Khweek et al., 2013). In contrast, a more recent study demonstrated that in J77A.1 macrophage-like cells, p62 as well as other autophagy adaptors are found on less than 10% of LCVs up to 9 hours post-infection (Omotade and Roy, 2020). As the LCV membrane undergoes remodeling throughout the course of infection, it is plausible that association with autophagic markers is a dynamic process coordinated by different effectors at distinct stages of infection. Identification of effectors that influence autophagy will lead to a more comprehensive understanding of how *L. pneumophila* exploits this highly conserved pathway during host infection.

In this study, we present *L. pneumophila* effector Lpg2411 as an autophagy-stimulating protein that accumulates in the host cell late during infection. We show that this effector localizes to autophagosomes, associates with ubiquitinated proteins, co-localizes with p62, and contributes to enhancing autophagy. Given that *L. pneumophila* encodes several autophagy-inhibiting effectors to avoid autophagosomal degradation, our finding of an autophagy-promoting effector supports the idea that *L. pneumophila* may employ a coordinated mechanism to exploit this pathway.

## Results and Discussion

### Lpg2411 is a membrane-associated protein that binds PI(3)P

Lpg2411 has previously been shown to be a Dot/Icm substrate, but its function has not yet been characterized (Burstein et al., 2009). We first investigated the subcellular localization of mCherry-Lpg2411. Confocal microscopy of HeLa cells transiently producing mCherry-Lpg2411 displayed two different populations of cells with distinct localization patterns: a pattern of foci or punctae dispersed throughout the cell (“punctae”) or a collection of tubular structures (“tubules”) (Fig. 1A). Quantification of confocal micrographs revealed that 54 ± 13% of transiently transfected HeLa cells displayed mCherry-Lpg2411 as punctae and 46 ± 8% of HeLa cells displayed mCherry-Lpg2411 as tubules (Fig. 1A). We reasoned that the punctate localization could indicate that Lpg2411 accumulates on membrane-bound compartments. Immunogold transmission electron microscopy of CHO cells ectopically producing GFP-Lpg2411 revealed localization of Lpg2411 on vesicular compartments as well as elongated structures (Fig. S1A). This led us to speculate that Lpg2411 may interact with a membrane-associated component such as a protein or lipid.

**Figure 1:**
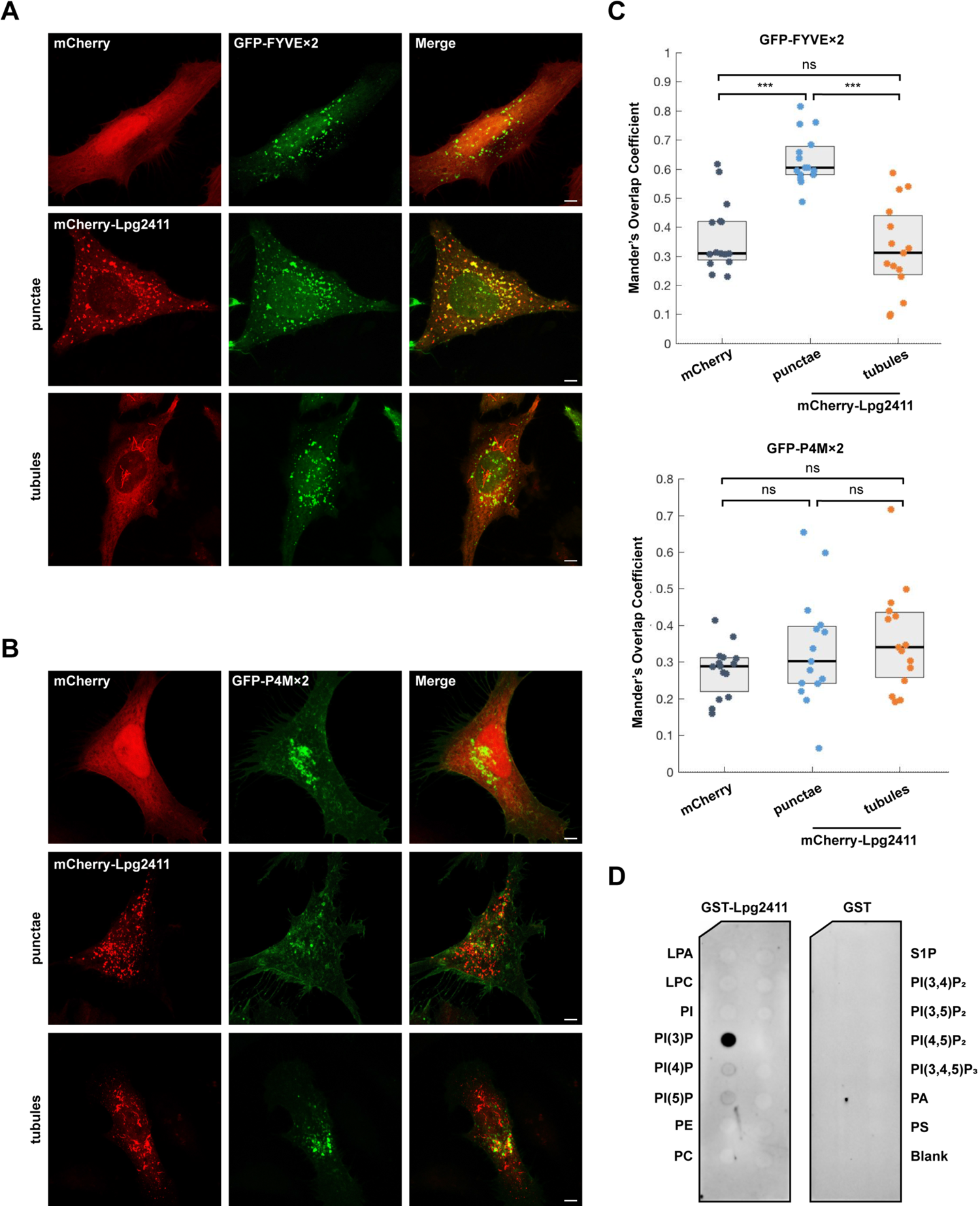
Lpg2411 localizes to PI(3)P-positive membranes. (A) Confocal images of HeLa cells transiently co-transfected with plasmids encoding mCherry-Lpg2411 and either GFP or GFP-2×FYVE. Scale bar, 5 µm. (B) Confocal images of HeLa cells transiently co-transfected with plasmids encoding mCherry-Lpg2411 and GFP or GFP-P4M×2. Scale bar, 5 µm. (C) Mander’s Overlap Coefficient for fluorescence signals of EGFP-tagged and mCherry-tagged proteins, as specified. The plot shows the median (black vertical line) and interquartile range (25-75) (gray box) from 15 different cells for each condition. Individual dots represent the Mander’s Overlap Coefficient obtained from a single cell; ***, p ≤ 0.001. (D) Protein–lipid overlay assay displays that GST-Lpg2411 specifically binds PI(3)P. A membrane spotted with each lipid species was incubated with purified GST-Lpg2411. GST-Lpg2411 that bound to the membrane was identified via detection with anti-GST and an HRP-conjugated secondary antibody. LPA, lysophosphatidic acid; LPC, lysophosphocholine; PI, phosphatidylinositol; PE, phosphatidylethanolamine; PC, phosphatidylcholine; S1P, sphingosine-1-phosphate; PI, phosphatidylinositol; P, phosphate; P_2_, biphosphate; P_3_, triphosphate; PA, phosphatidic acid; PS, phosphatidylserine.

Several previously identified effectors recognize membrane compartments by selectively binding phosphoinositide lipids (PIP) (reviewed in (Pike et al., 2019a)). Therefore, we next asked whether Lpg2411 preferentially localizes to membranes enriched in either phosphatidylinositol-3-phosphate (PI(3)P) or phosphatidylinositol-4-phosphate (PI(4)P). A growing number of *L. pneumophila* effectors preferentially bind PI(3)P and PI(4)P and these PIPs are enriched on the LCV membrane at different times throughout infection (Weber et al., 2018; Weber et al., 2014). Using confocal microscopy, we visualized and quantified the subcellular distribution of mCherry-Lpg2411 relative to the PI(3)P marker GFP-FYVE×2 (Hammond and Balla, 2015) or the PI(4)P marker GFP-P4M×2 (Hammond et al., 2014). Co-localization was quantified by calculating the Mander’s Overlap Coefficient (MOC; where a value of 1 represents complete overlap and 0 represents random localization) as performed previously (Manders et al., 1993; Pike et al., 2019b). This quantification indicated that the Lpg2411 punctae were PI(3)P-positive whereas Lpg2411 tubules were not (MOC= 0.63 ± 0.02, 0.32 ± 0.04, respectively; Fig. 1A, B). The MOC of GFP-FYVE×2 and mCherry-Lpg2411 punctae was significantly higher than the MOC of mCherry alone and GFP-FYVE×2 (MOC= 0.36 ± 0.02; p<0.0001, Kruskal-Wallis test with Dunn-Sidak post-hoc test), whereas the MOC of GFP-FYVE×2 and mCherry-Lpg2411 tubules was not significantly different from the MOC of mCherry alone and GFP-FYVE×2 (p>0.9999, Kruskal-Wallis test with Dunn-Sidak post-hoc test) (Fig. 1C). Neither the Lpg2411 punctae, nor the Lpg2411 tubules co-localized with the PI(4)P marker GFP-P4M×2 (MOC= 0.33 ± 0.04, 0.36 ± 0.04, respectively; Fig. 1B, C), as these MOCs were not significantly different than the MOC of mCherry alone and GFP-P4M×2 (MOC= 0.27 ± 0.02, p>0.9999, Kruskal-Wallis test with Dunn-Sidak post-hoc test) (Fig. 1C). These results suggest that the punctae Lpg2411 are PI(3)P-positive compartments.

To determine if Lpg2411 directly binds PI(3)P, we performed an *in vitro* protein-lipid overlay assay using a purified GST-fusion protein (Fig. S1B). After purification, GST-Lpg2411 was incubated with a nitrocellulose membrane pre-spotted with seven PIPs and several other biologically relevant lipids. GST-Lpg2411 predominantly bound PI(3)P, and weaker signals were detected for PI(4)P and PI(5)P (Fig. 1D). Taken together, these findings indicate that Lpg2411 directly binds PI(3)P *in vitro* and is present on PI(3)P-positive membranes in HeLa cells. Whether Lpg2411 requires PI(3)P for membrane recognition and attachment is unclear. Treatment with the PI3-kinase inhibitor, wortmannin, abolished the tubular localization of mCherry-Lpg2411 but did not disrupt the punctae localization of mCherry-Lpg2411 in HeLa cells, suggesting that tubule formation or stability may require the presence PI(3)P (data not shown).

### Lpg2411 selectively localizes to autophagosomes and co-localizes with the autophagic receptor, p62

PI(3)P is predominantly enriched on early and late endosomal membranes as well as autophagosomal compartments (Simonsen and Tooze, 2009; Wurmser et al., 1999). To further probe the function of Lpg2411, we next sought to determine the identity of the PI(3)P-positive compartments recognized by Lpg2411. We found that in transiently transfected HeLa cells, mCherry-Lpg2411 localized to compartments containing the autophagosome marker, GFP-LC3 (Fig. 2A). Both Lpg2411-positive punctae as well as short tubules co-localized with GFP-LC3. We did not observe co-localization between mCherry-Lpg2411 and markers of early endosomes (GFP-Rab5) or late endosomes/lysosomes (LAMP1-GFP), consistent with the notion that Lpg2411 selectively localizes to autophagosomes.

**Figure 2:**
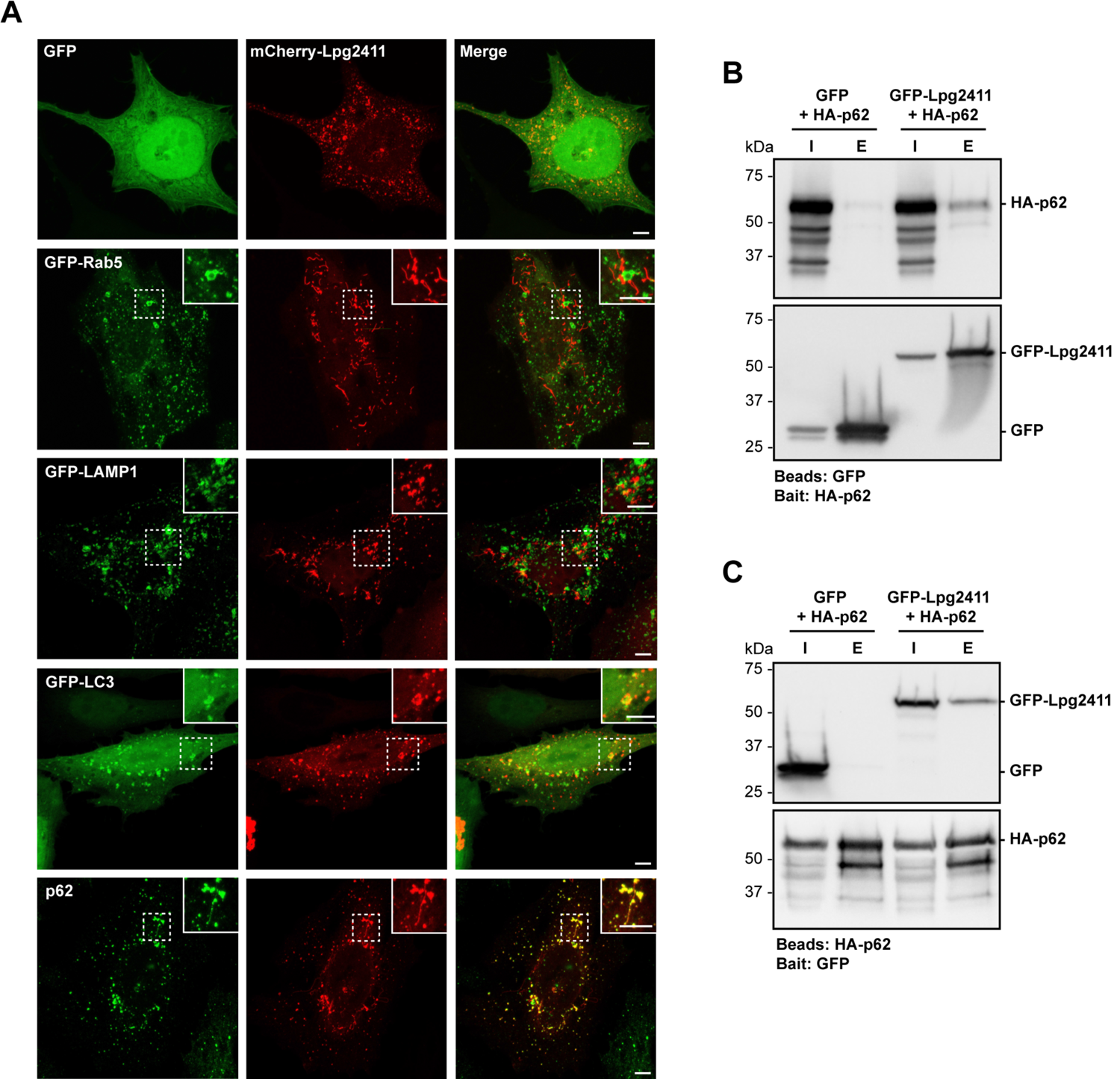
Lpg2411 localizes to autophagosomes and binds an autophagic receptor. (A) Confocal images of HeLa cells transiently co-transfected with mCherry-Lpg2411 and GFP, GFP-Rab5, GFP-LAMP1, or GFP-LC3, or singly transfected with mCherry-Lpg2411 and immunostained for endogenous p62 with anti-p62 followed by Alexa Fluor 488-conjugated secondary antibody. Scale bar, 5 µm. (B) Lysates from HEK293T cells co-transfected with HA-p62 and GFP or GFP-Lpg2411 subjected to a co-immunoprecipitation assay using anti-HA magnetic beads. Cleared lysates, or inputs (I), and elutions (E) from the assay separated via SDS-PAGE and transferred to a PVDF membrane. Western blots were probed with HRP-conjugated anti-GFP and anti-HA. HA-p62 = 63 kDa, GFP= 27 kDa, GFP-2411= 59 kDa. (C) Reciprocal co-immunoprecipitation performed using lysates from HEK293T cells co-transfected with HA-p62 and GFP or GFP-Lpg2411 and GFP-Trap magnetic agarose beads. Resulting western blots were probed with anti-HA and HRP-conjugated anti-GFP.

Given this observation, we reasoned that in addition to binding PI(3)P, Lpg2411 may interact with another component that facilitates selective binding to autophagosomes. We therefore probed whether Lpg2411 associates with proteins involved in autophagy. The autophagic receptor, p62/SQSTM1, localizes to nascent autophagosome membranes and serves as an adaptor between autophagosomes and ubiquitinated proteins destined for autophagic degradation (Lamark et al., 2009; Pankiv et al., 2007). HeLa cells ectopically producing mCherry-Lpg2411 were immunostained with a p62 antibody and visualized with confocal microscopy. Indeed, we observed co-localization between mCherry-Lpg2411 and endogenous p62 (Fig. 2A). To determine whether Lpg2411 associates with p62, we performed co-immunoprecipitation assays. Transiently transfected HEK293T cells ectopically producing hemagglutinin (HA) tagged p62 and GFP-Lpg2411 or GFP alone were lysed and GFP was immunoprecipitated using GFP-Trap magnetic agarose beads. GFP-Lpg2411, but not GFP alone, precipitated HA-p62 from the cellular lysate (Fig. 2B). In a reciprocal co-immunoprecipitation assay, HA-p62 bound to anti-HA magnetic beads precipitated GFP-Lpg2411, but not GFP, from cellular lysate (Fig. 2C). These data provide evidence that Lpg2411 associates (directly or indirectly) with the autophagic receptor, p62. This, in turn, could facilitate Lpg2411’s ability to selectively localize to autophagosomes.

### Lpg2411 associates with ubiquitinated proteins

p62 specifically links ubiquitinated proteins to autophagosomes by interacting with LC3 on the autophagosome membrane, leading to the encapsulation of these ubiquitinated proteins in the double membrane compartment (Lamark et al., 2009; Pankiv et al., 2007). We therefore postulated that Lpg2411 may also interact with the p62-ubiquitinated protein complex. Confocal microscopy revealed that mCherry-Lpg2411 co-localized with HA-Ubiquitin on punctate structures in transiently transfected HeLa cells, indicating that this effector and ubiquitinated proteins are in close proximity (Fig. 3A). To further probe this association, we assessed whether Lpg2411 forms a complex with ubiquitinated proteins. HEK293T cells producing mCherry-Lpg2411 or mCherry alone were lysed and mCherry was immunoprecipitated from the lysate via an RFP-Trap assay. Via immunoblot, we found that mCherry-Lpg2411, but not mCherry alone precipitated high molecular weight ubiquitin species (Fig. 3B). These data support the notion that Lpg2411 directly or indirectly interacts with ubiquitinated proteins.

**Figure 3:**
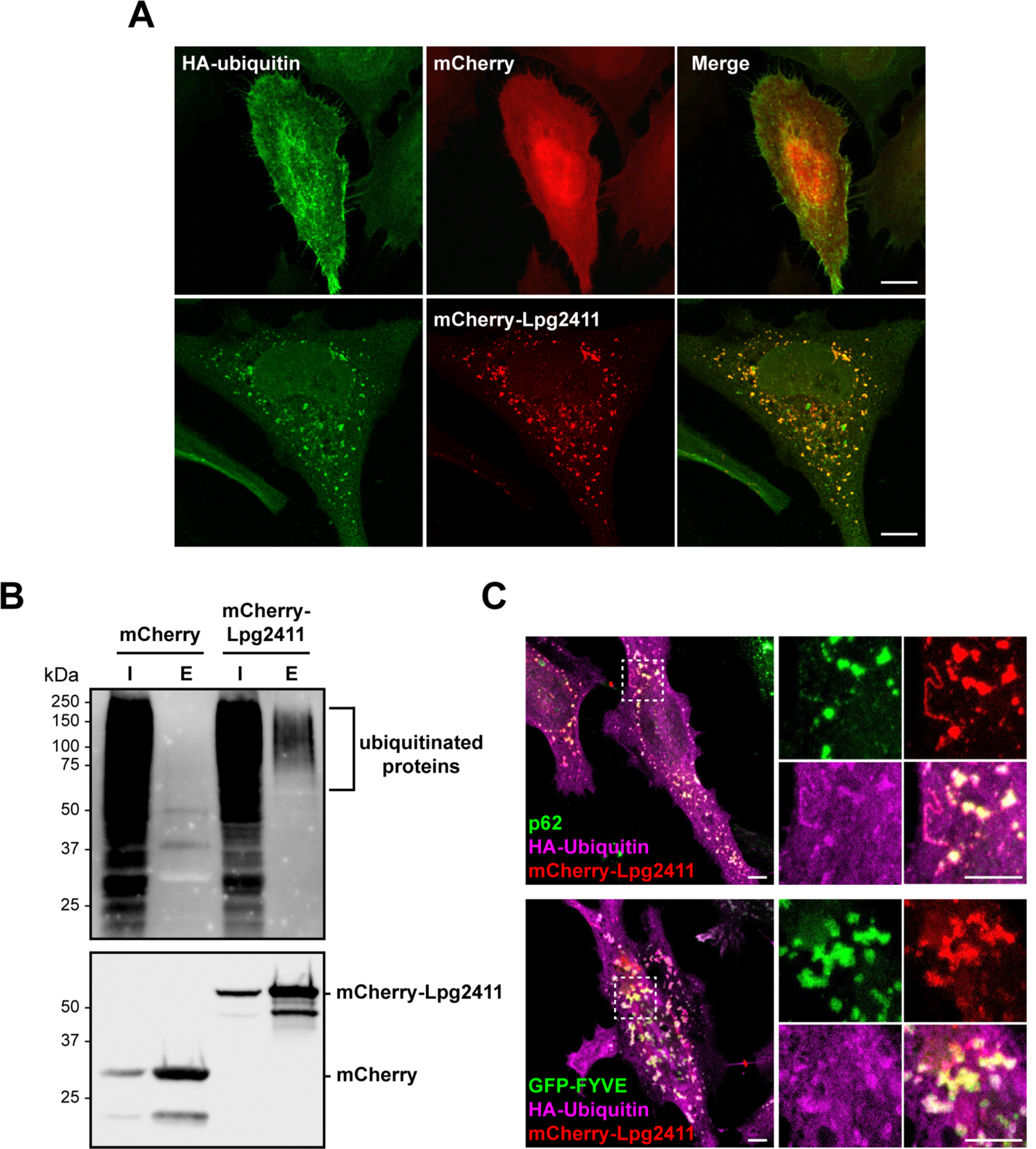
Lpg2411 binds ubiquitinated proteins on autophagosomes. (A) Confocal images of HeLa cells co-expressing HA-tagged ubiquitin and mCherry-Lpg2411 and immunostained with anti-HA followed by Alexa Fluor 488-conjugated anti-rat. Scale bar, 5 µm. (B) Confocal images of HeLa cells expressing HA-tagged ubiquitin, mCherry-Lpg2411, and GFP-2×FYVE were immunostained with anti-HA followed by Alexa Fluor 647-conjugated anti-rat. (C) HeLa cells co-expressing HA-Ubiquitin and mCherry-Lpg2411 were immunostained with anti-p62 and anti-HA followed by Alexa Fluor 488-conjugated and Alexa Fluor 647-conjugated secondary antibodies, respectively. Scale bar, 5 µm. (D) HEK293T cells expressing mCherry or mCherry-Lpg2411 were lysed and immunoprecipitated with RFP-Trap magnetic agarose beads. Cleared lysates, or inputs (I), and elutions (E) from the assay separated via SDS-PAGE and transferred to a PVDF membrane were probed with anti-ubiquitin and anti-mCherry. mCherry= 29 kDa, mCherry-2411= 61 kDa.

We next investigated whether the ubiquitinated proteins associating with Lpg2411 were targeted for autophagy by examining localization between these components and endogenous p62. In HeLa cells transiently transfected with mCherry-Lpg2411 and HA-Ubiquitin, and immunostained with antibodies against HA and p62, we observed areas of overlap between mCherry-Lpg2411, HA-Ubiquitin, and p62 (Fig. 3C). Thus, Lpg2411 was present at sites of autophagic degradation of ubiquitinated substrates. We also observed that mCherry-Lpg2411 and HA-Ubiquitin co-localized at PI(3)P-positive sites marked by GFP-FYVE×2 (Fig. 3C), further suggesting that this complex is present on autophagosomes.

### Lpg2411 accumulates in the host cell late during infection

Our data provide strong evidence that Lpg2411 targets autophagosomes and suggest that Lpg2411 may modulate autophagy. Several previous studies have identified *L. pneumophila* effectors that interfere with autophagy during the early hours of host infection (Arasaki et al., 2017; Choy et al., 2012). To determine when Lpg2411 is translocated during infection, U937 macrophages were infected with a Δ*lpg2411* strain carrying the pMMB207c-HA×4-*lpg2411* plasmid (Δ*lpg2411* + pMMB207c-HA×4-*lpg2411*) at an MOI of 20 (Fig. 4A). At 1, 5, 10, 14 and 18 hours post-infection, cells were immunolabeled with HA and *L. pneumophila* antibodies and the number of cells displaying HA signal was quantified. Although no signal was detected at 1-5 hours post-infection, at 10 hours, 51% of the cells displayed HA-Lpg2411 signal, and the percentage continued to rise to 66% at 14 hours, and 73% at 18 hours post-infection. While the lack of HA-Lpg2411 signal in the early hours of infection could indicate that this effector protein is not translocated until later during infection, it is also possible that early during infection Lpg2411 is translocated at levels below our detection limit and accumulates to higher, more readily detectable levels as the infection progresses.

**Figure 4:**
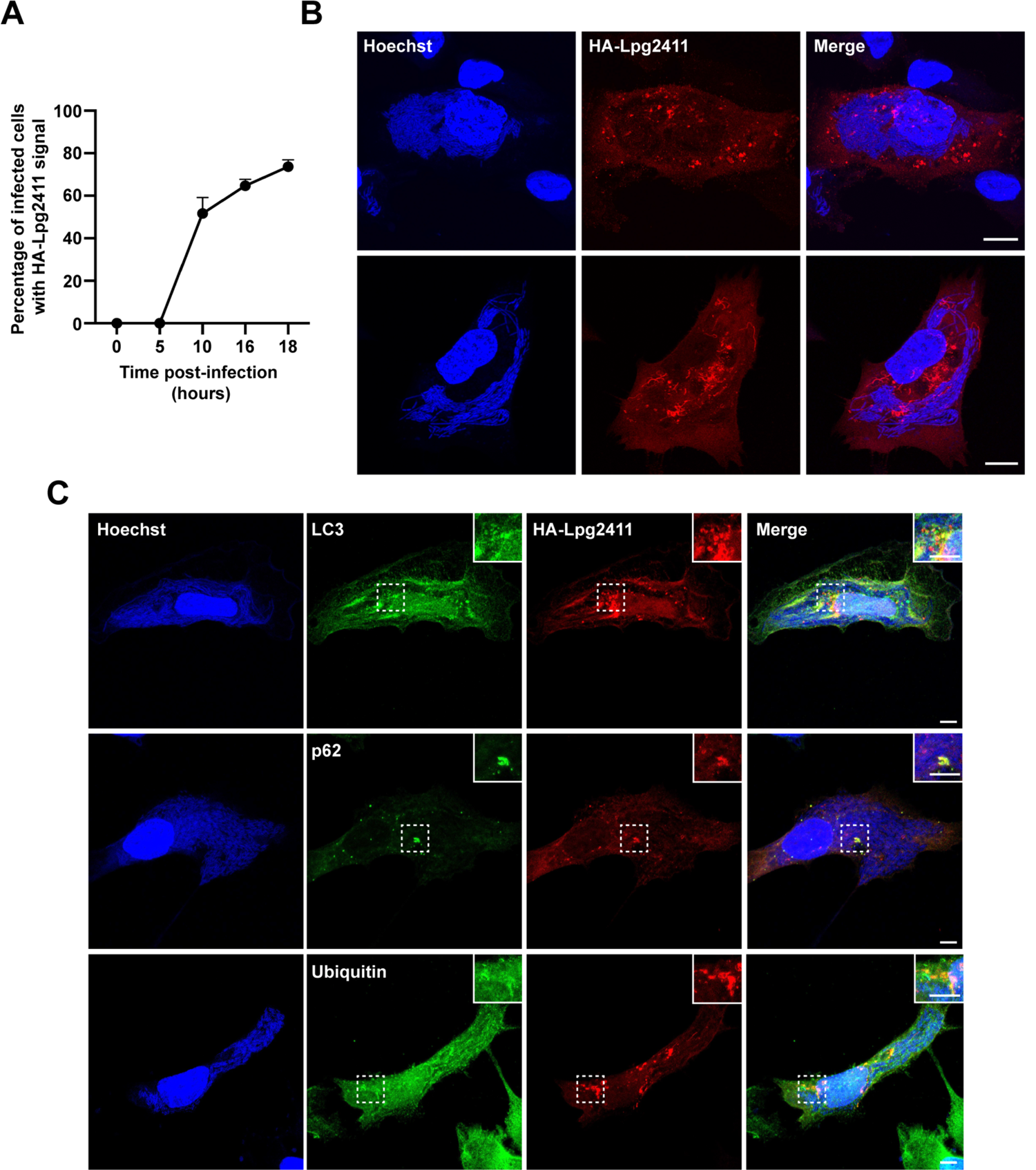
Lpg2411 is translocated during late stages of infection and localizes to autophagosomes. (A) U937 cells were infected with Δ*lpg2411* + pMMB207c-HA×4-*lpg2411* for indicated time points before immunostaining for HA-Lpg2411 with anti-HA and TexasRed-conjugated secondary antibody. Infected cells harboring translocated HA-Lpg2411 were counted and compared to the total infected cells. (B) HT1080 cells infected with Δ*lpg2411* pMMB207c-HA×4-*lpg2411* for 15 hours were immunostained for HA-Lpg2411 with anti-HA and TexasRed-conjugated secondary antibody. Hoechst was used to visualize the bacteria and host nucleus. (C) HT1080 cells were infected as in B and immunostained for anti-HA with TexasRed-conjugated secondary antibody and either anti-LC3, -p62, or -FK2 with AlexaFluor 488-conjugated secondary antibodies. Hoechst was used to visualize the bacteria and host nucleus.

### Lpg2411 is dispensable for *L. pneumophila* intracellular growth in U937 macrophages and HT1080 cells

To address whether Lpg2411 impacts the intracellular survival of *L. pneumophila*, we employed HT1080 human fibrosarcoma cells as a host model. HT1080 cells undergo increased membrane ruffling and endocytosis due to a constitutively active N-Ras allele (Donaldson, 2019; Williamson and Donaldson, 2019), resulting in efficient bacterial uptake. Once *L. pneumophila* entered HT1080 cells, they grew robustly (Fig. S2A); and as expected, growth of the Dot/Icm-defective variant was efficiently restricted (Fig. S2B). Wild-type vacuoles displayed features described during infection of professional phagocytes, including decoration with ubiquitin (Fig. S2D, E); and co-localization with organelle markers such as YFP-KDEL, Calnexin (ER) and Rab1 (Golgi) (Fig. S2F, G). Therefore, HT1080 cells can serve as an effective cell line to study *L. pneumophila* infection.

We performed intracellular replication assays in both U937 and HT1080 cells. Host cells were infected with either wild-type, Δ*lpg2411* + pMMB207c-HA×4, Δ*lpg2411* + pMMB207c-HA×4-*lpg2411*, or the translocation deficient Δ*dotA* strain and intracellular growth was compared between strains at 48 and 72 hours post-infection. We observed that in HT1080 cells, the wild-type strain was able to establish intracellular replication leading to high numbers of bacteria comparable to infection in U937 (Fig. S3B). The avirulent strain lacking a functional Dot/Icm T4SS was unable to replicate intracellularly within the HT1080 cell line, as observed in U937 cells. Additionally, a strain lacking *lpg2411* carrying an empty plasmid (Δ*lpg2411* + pMMB207c-HA×4) as well as a complemented strain (Δ*lpg2411* + pMMB207c-HA×4-*lpg2411*) survived intracellularly comparably to wild-type in both HT1080 and U937 cells, suggesting that Lpg2411 is dispensable for infection. A significant growth defect may not have been observed due to the functional redundancy among *L. pneumophila* effectors (Ghosh & O’Connor, 2017). It is possible that another effector has a similar function as Lpg2411, in which case a phenotype caused by the absence of Lpg2411 would be masked in this assay.

### Lpg2411 localizes to autophagosomes late during infection of HT1080 cells

Since Dot/Icm-mediated translocation of Lpg2411 was detected late during infection, we next asked whether Lpg2411 binds autophagosomes at this stage of infection. To address this question, we first visualized the subcellular localization of translocated HA-Lpg2411 in HT1080 cells infected with Δ*lpg2411* + pMMB207c-HA×4-*lpg2411* for 15 hours. Similar to our observations in transiently transfected HeLa cells, HA-Lpg2411 localized to both punctate and tubular structures within the infected host cell (Fig. 4B). Using antibodies directed against various organelle markers (Fig. 4C), our confocal microscopy analyses revealed that HA-Lpg2411 co-localized with LC3, p62, and ubiquitin in infected HT1080 cells at 15 hours post-infection. These findings corroborated our results from Figures 2 and 3 that Lpg2411 is present on autophagosomes that contain p62 and ubiquitin.

We observed that ubiquitin was present on the LCV throughout infection, similar to previous findings (Dorer et al., 2006). Intriguingly, although Lpg2411 co-localized with ubiquitin in transiently transfected HeLa cells, we did not observe Lpg2411 on the LCV membrane at 15 hours post-infection. Notably, p62 was not observed on or around the LCV either, consistent with a previous report’s finding that autophagy receptors are excluded from the LCV membrane (Omotade and Roy, 2020). We speculate that autophagy receptors, like p62, are not recruited to ubiquitin on the LCV due to the unique ubiquitin linkages catalyzed by members of the SidE family of effectors. These effectors ligate phosphoribosylated ubiquitin and it is possible that this modification remains unrecognized by autophagic receptors and Lpg2411 (Bhogaraju et al., 2016; Qiu et al., 2016). Alternatively, it is conceivable that Lpg2411 does not bind ubiquitin directly and that it instead more closely interacts with p62, which is excluded from the vacuole.

### Lpg2411 stimulates autophagy late during infection

Due to the association of Lpg2411 with autophagosomes, we next investigated whether Lpg2411 interferes with regulation of autophagy. To study this cellular process, the levels of lipidated LC3 (LC3-II), a marker of autophagy stimulation, were quantified in the presence or absence of Bafilomycin A1, as done previously (Choy et al., 2012). This chemical compound inhibits fusion of autophagosomes with lysosomes thereby preventing degradation of LC3-II. Using this experimental principle, HT1080 cells were infected with wild-type, Δ*lpg2411* + pMMB207c-HA×4 (empty vector, EV), or Δ*lpg2411* + pMMB207c-HA×4-*lpg2411* (HA-Lpg2411) strains for 15 hours and either treated with Bafilomycin A1 or left untreated for 2 hours. Cells were then lysed and the lysates were separated via SDS-PAGE and transferred to a PVDF membrane for detection of LC3 by western blotting (Fig. 5A). The intensity signals of LC3-II, as well as actin as a loading control, were quantified using ImageJ. LC3-II signal intensity was normalized to the actin signal and this value is represented as a value relative to uninfected cells. Cells infected with the wild-type strain and treated with Bafilomycin A1 showed significantly less LC3-II compared to uninfected cells, (p=0.0018; data not shown), indicating that overall *L. pneumophila* inhibits autophagy during late stages of infection, in agreement with previous data (Rolando et al., 2016). However, compared to wild-type, cells infected with Δ*lpg2411* + pMMB207c-HA×4 had significantly less LC3-II (p=0.0475, One-way ANOVA with Tukey’s multiple comparisons test), suggesting that the strain lacking Lpg2411 results in significantly less stimulation of autophagy during infection (Fig. 5B). In cells infected with a complemented strain (Δ*lpg2411* + pMMB207c-HA×4-*lpg2411*), LC3-II levels were comparable to cells challenged with wild-type (p=0.9968, One-way ANOVA with Tukey’s multiple comparisons test) (Fig. 5B). In turn, a significant difference was observed between cells infected with Δ*lpg2411* + pMMB207c-HA×4 and cells infected with the complemented strain (p=0.0219, One-way ANOVA with Tukey’s multiple comparisons test). These data indicate that Lpg2411 is contributing to the stimulation of autophagy during *L. pneumophila* infection.

**Figure 5:**
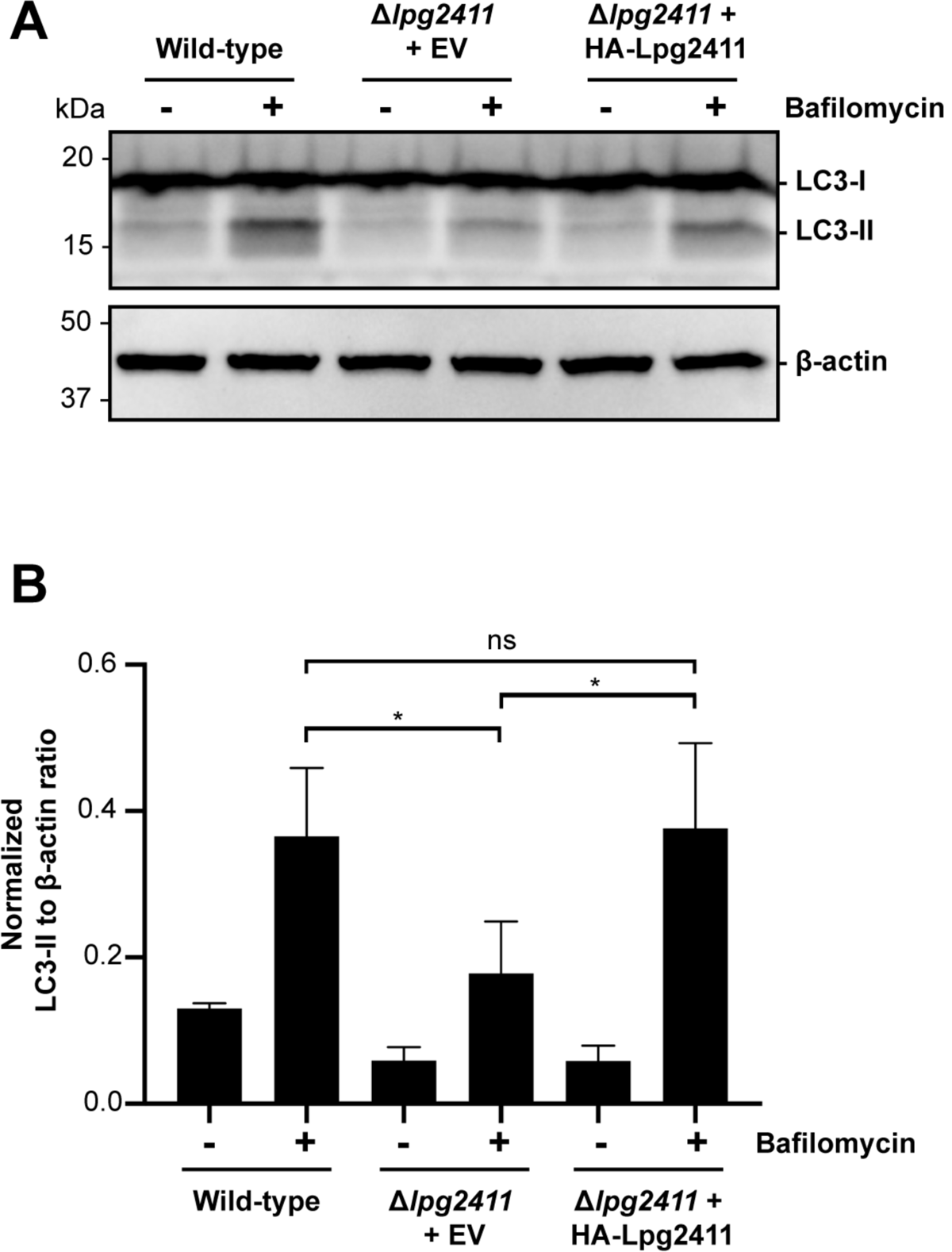
Lpg2411 stimulates autophagy late during infection. (A) HT1080 cells were infected with wild-type, Δ*lpg2411* + pMMB207c-HA×4 (empty vector, EV), Δ*lpg2411* + pMMB207c-HA×4-*lpg2411* (HA-Lpg2411) for 15 hours. Cells were then left untreated or treated with 160 nM Bafilomycin A1 for 2 hours before lysing and separating via SDS-PAGE. Proteins were transferred to a PVDF membrane and probed with anti-LC3 and anti-actin. LC3-I= 19 kDa, LC3-II= 17 kDa, actin= 42 kDa. (B) LC3-II bands were quantified for each condition and normalized to the signal for actin using ImageJ. Normalized values are displayed relative to uninfected cells. n=3; *, p<0.05.

### Autophagy serves two main roles in a eukaryotic cell

The first is to clear damaged organelles, protein aggregates, or invading pathogens. The second is to liberate nutrients for the cell during periods of starvation by breaking down cellular materials. Autophagosomes can engulf and degrade pathogens by fusion with lysosomes. It would therefore be beneficial for *L. pneumophila* to inhibit this degradative process at first to ensure intracellular survival. Indeed, effector-mediated inhibition of autophagic processes has previously been documented during *L. pneumophila* infection. Effector proteins RavZ, Lpg1137, LegS2/LpSpI, and Lpg2936 were shown to inhibit autophagy during *L. pneumophila* infection. RavZ, capable of deconjugating Atg8 from phosphatidylethanolamine, inhibits autophagosome maturation at around 2 hours post-infection (Choy et al., 2012; Horenkamp et al., 2015). Lpg1137 cleaves Syntaxin17, a SNARE important for autophagosome maturation, for up to 8 hours post-infection (Arasaki et al., 2017). Thus, autophagy is actively suppressed during the early stages of *L. pneumophila* infection. Subsequently, between 8 and 20 hours of infection, LegS2/LpSpl depletes the cell of sphingosine-1-phosphate leading to an overall decrease in autophagosome formation during the later stages of infection (Rolando et al., 2016). Lpg2936 was found to target autophagy at the transcription level, modifying the promoter regions of autophagy related genes and decreasing their expression (Abd El Maksoud et al., 2020).

Thus, *L. pneumophila* employs multiple strategies to inhibit autophagy possibly to prevent its elimination via autophagolysosome degradation. However, if inhibiting autophagy was the sole objective during *L. pneumophila* infection, inhibitors of autophagy would theoretically stimulate intracellular survival. Interestingly, this is not always the case. Studies have shown that inhibiting autophagy with the compound 3-methyladenine led to an increase in bacterial degradation (Amer and Swanson, 2005) and activating autophagy through amino acid starvation increased intracellular replication (Swanson and Isberg, 1995), suggesting that aspects of autophagy are beneficial for *L. pneumophila* infection. Conversely, another study showed that activation of autophagy decreased replication and inhibition increased bacterial replication (Matsuda et al., 2009). While inhibiting autophagy may be advantageous to *L. pneumophila* in certain aspects, these findings suggest that there may be a fine balance of autophagic processes that must be maintained. Not only does autophagy rid the cell of invading pathogens, but it contributes to the cellular homeostasis by eliminating dysfunctional organelles and protein aggregates that could cause harm to the cell. In turn, the breakdown of these macromolecules releases the building blocks and nutrients necessary to generate new structures. We speculate that stimulation of autophagy by effectors like Lpg2411 may counterbalance the detrimental effects of infection and support cellular homeostasis, which, in turn, would benefit *L. pneumophila* intracellular survival.

Alternatively, as the LCV grows and the cell becomes depleted of nutrients, *L. pneumophila* may activate autophagy to gain access to nutrients during late stages of infection. The aforementioned breakdown of dysfunctional organelles or protein aggregates releases free amino acids, a key carbon source for intracellular *L. pneumophila* (Tesh et al., 1983). Altogether, *L. pneumophila* actively inhibits autophagy for the escape from the degradative autophagosome-lysosome pathway. The emergence of autophagy-stimulating effectors, such as Lpg2411, could suggest that *L. pneumophila* counterbalances the autophagic response that also allows for cellular homeostasis and nutrient acquisition.

## Materials and methods

### Strains, media, and reagents

All the strains used in this study are listed in Table S1. *L. pneumophila* strain Lp01 (*hsdR rpsL*) and Lp01Δ*dotA* (T4SS-) are strains derived from *L. pneumophila* strain Philadelphia-1 (Berger and Isberg, 1993) and were cultured as previously described (Morris et al., 1979). HEK293T cells (ATCC CRL-1573) and HT1080 human fibrosarcoma cells (ATCC CCL-121) were cultured in DMEM supplemented with 2 mM L-glutamine and 10% FBS, and incubated in 5% CO_2_ at 37°C. HeLa cells (ATCC CCL-2) and U937 monocytes (ATCC CRL-1593.2) were cultured in RPMI 1640 supplemented with 2 mM L-glutamine and 10% FBS, and incubated as above. U937 monocytes were differentiated by supplementing the media with 10 ng/mL 12-O-tetradecanoylphorbol-13-acetate. Bafilomycin A1 Ready Made Solution was purchased from Sigma-Aldrich (SML1661). Antibodies were purchased from ThermoFisher Scientific (goat anti-rabbit Alexa Fluor 568, A-11036; goat anti-rabbit Alexa Fluor 488, A-11008; goat anti-mouse Alexa Fluor 488, A-11001; goat anti-rat Texas Red-X, T-6392; goat anti-rat Alexa Fluor 647, A-21247; monoclonal mouse anti-GST, MA4-004; polyclonal rabbit anti-mCherry, Pa5-34974; polyclonal rabbit anti-ubiquitin, 10201-2-AP; HRP-conjugated anti-mouse antibody, NA931; HRP-conjugated anti-rabbit antibody, 31460; HRP-conjugated anti-rat antibody 31470; HRP-conjugated streptavidin antibody, SA10001), Sigma-Aldrich (monoclonal rat anti-HA, 11867423001A; polyclonal rabbit anti-ICDH, ABS2090; monoclonal mouse anti-β actin, A2228), Abcam (HRP-conjugated anti-GFP, ab6663), Novus Biologicals (polyclonal rabbit anti-LC3B, NB100-2220), Cell Signaling Technologies (monoclonal mouse anti-p62, 88588), and Enzo Life Sciences (monoclonal mouse anti-mono-and polyubiquitinylated conjugates, FK2, BML-PW0150).

### Construction of expression clones

Bacterial strains, plasmids and oligonucleotides used in this study are listed in Table S1 and Table S2. *lpg2411* was cloned into the destination vectors, pDEST15, 362 pCS Cherry DEST and pcDNA6.2/N-EmGFP-DEST via Gateway^TM^ cloning technology to generate translational fusions with either GST, GFP or mCherry, respectively. The plasmid pMMB207c-HA×4 carrying a hemagglutinin tag under an IPTG inducible promoter was a kind gift from Dr. Gunnar Schroeder (Queen’s University Belfast). The pMMB207c-HA×4-*lpg2411* plasmid was generated by cloning *lpg2411* into pMMB207c-HA×4 at the BamHI and SalI restriction sites. The 362 pCS Cherry DEST plasmid was a gift from Nathan Lawson (Addgene plasmid # 13075) (Villefranc et al., 2007).

### Construction of the *L. pneumophila* Δ*lpg2411* deletion mutant

An in-frame deletion of *lpg2411* was obtained by allelic exchange using the pNTPS138 plasmid that carries a chloramphenicol resistance cassette and the *sacB* gene (sucrose sensitivity). The pNTPS138 plasmid was a gift from Howard Steinman (Addgene plasmid #41891); ∼500 nucleotide-fragments flanking *lpg2411* were PCR amplified and cloned into pNTPS138 at the HindIII and EcoRI restriction sites. This plasmid was then transformed into the Lp01 strain by electroporation as previously described (Chen et al., 2006) and single recombinants were selected on CYE agar plates containing chloramphenicol. Selected colonies were then plated on CYE agar with 8% sucrose, and sucrose selection was used to identify colonies in which the plasmid was lost. Deletion of the wild-type allele was verified by PCR analysis and confirmed by sequencing.

### Recombinant protein production and purification

Lpg2411 was produced as a GST fusion protein in *E. coli* BL21 (DE3) at 25°C overnight after induction with 0.5 mM isopropyl-β-dithiogalactopyranoside (IPTG). *E. coli* cells producing GST-Lpg2411 were harvested and resuspended in PBS supplemented with 1 mM MgCl_2_ and 1 mM β-mercaptoethanol (PBS-MM) followed by lysis using the LV10 microfluidizer (Microfluidics). Cell lysate was centrifuged at 24,000 × g for 35 min, and the supernatant was incubated with pre-equilibrated glutathione sepharose 4B (GE Healthcare) for 2 hours at 4°C. The resin was washed three times with PBS-MM, and proteins were eluted in 50 mM Tris-HCl (pH 8) containing 10 mM reduced glutathione (Sigma). Glutathione was removed using a desalting Zeba column following manufacturer instructions (ThermoFisher Scientific).

### Protein-lipid overlay assay

Protein-lipid overlay assays were performed using commercially available PIP strips (Echelon Biosciences Inc.). Nitrocellulose membranes pre-spotted with different phospholipids were blocked with 2% nonfat milk in PBST [PBS and 0.1% Tween-20 (v/v) pH 7.5] for 1 hour at room temperature. The blocked membranes were incubated with purified GST-Lpg2411 or GST alone (0.8 µg/mL in blocking buffer) overnight at 4°C. Binding of the GST-fusion protein to lipids was visualized with a mouse anti-GST antibody (1:2000) and an HRP-conjugated anti-mouse antibody (1:20,000).

### Confocal microscopy

Constructs based on pEGFP-C1 or 362 pCS mCherry DEST as listed in Table S1 were transiently transfected into HeLa cells for 16-20 hours using Lipofectamine 3000 transfection reagent (ThermoFisher Scientific). Cells were then fixed in PBS with 4% paraformaldehyde for 20 min at room temperature and coverslips were mounted using ProLong diamond anti-fade mountant (ThermoFisher Scientific). For immunostaining experiments, fixed cells were permeabilized with 0.1% saponin incorporated in all staining steps or 0.1% Triton X-100. Fixed cells were incubated in blocking buffer (10% goat serum in PBS) for 1 hour, incubated with primary antibodies in blocking buffer for 1 hour and finally incubated with secondary antibodies in blocking buffer for 1 hour. Confocal imaging was performed on a Zeiss LSM 880 laser-scanning confocal microscope using a 63× Plan-Apochromat objective lens (numerical aperture of 1.4) and operated with ZEN software (Carl Zeiss, Inc).

### Colocalization quantification and statistical analysis

For each condition, we acquired confocal images of 15 cells. Quantitative co-localization analysis was performed using the Volocity software (PerkinElmer, Waltham, MA) to calculate the Mander’s overlap coefficient (Manders et al., 1993) corresponding to the fraction of green voxels overlapping with red voxels in relation to the total green voxels. Normality of the distribution of variables was assessed using the Kolmogorov-Smirnov test and was then conformed visually. Data were non-parametrically distributed, and therefore to compare coefficients across the conditions tested, we used the Kruskal-Wallis test (H=18, p<1-13). The median and the interquartile range of coefficients were calculated for each condition. Dunn-Sidak post-hoc tests were conducted to statistically determine which groups differed while correcting for multiple comparisons. Statistical significance was set at p≤0.05. Matlab (Mathworks; Natick, MA; version 2016a) was used as statistical software.

### Immunogold transmission electron microscopy

#### Transmission electron microscopy

CHO cells transiently producing GFP-Lpg2411 were grown on 1.2 mm × 200 µm high pressure freezer carriers. Just prior to freezing, the medium was removed, and the carrier was filled with medium containing 20% BSA. Cells were frozen with a Leica EM Pact2 high-pressure freezer and then transferred under liquid nitrogen to a Leica AFS for freeze substitution. Samples were freeze-substituted in 0.1% uranyl acetate in 100% acetone at 90°C for 4–5 days. Samples were warmed to 45°C over 12 hours, washed with 100% acetone, and gradually infiltrated with increasing concentrations of Lowicryl HM20 monostep resin over a period of 2 days. The samples were embedded in Lowicryl HM20 Monostep resin and polymerized under UV light for 48 hours at 45°C and for an additional 48 hours at 23°C. The cells were sectioned using a Reichert–Jung Ultracut E ultramicrotome, and ultrathin sections were collected onto 200 mesh formvar/carbon-coated nickel grids.

#### Immunogold Labeling

Samples were blocked with 0.05 M glycine for 15 min and Aurion goat blocking solution for 30 min before being incubated on drops of anti-GFP antibody diluted to 2.8 µg/mL in Aurion 0.1% BSA-c 7.4 for 1 hour. Control grids were incubated on drops of ChromePure rabbit IgG (Jackson ImmunoResearch, Cat No. 011-000-003) diluted to 2.8 µg/mL in Aurion 0.1% BSA-c 7.4. Grids were washed on six drops of Aurion 0.1% BSA-c and incubated on drops of Aurion goat anti-rabbit IgG conjugated to 10 nm gold diluted 1:20 in Aurion 0.1% BSA-c for 2 hours. Grids were washed on six drops of Aurion 0.1% BSA-c, 3 drops of PBS, fixed on drops of 2% glutaraldehyde in PBS, and then washed on five drops of Nanopure water. The grids were then post-stained with 2% uranyl acetate in 50% methanol and Reynolds’ lead citrate. The samples were examined with a ZEISS Libra 120 transmission electron microscope operating at 120 kV, and images were acquired with a Gatan Ultrascan 1000 CCD camera.

#### Immunoprecipitation assays

For co-immunoprecipitation of Lpg2411 and p62, HEK293T cells were transfected with plasmids encoding HA-p62 and GFP-Lpg2411 or GFP alone using Lipofectamine 3000 reagent (ThermoFisher Scientific) according to the manufacturer’s instructions. GFP-Trap®_MA kit (Chromotek) or Pierce™ Anti-HA Magnetic Beads (ThermoFisher Scientific) were used to precipitate either GFP- or HA-tagged proteins, respectively. Transfected cells were lysed in lysis buffer (10 mM Tris/Cl pH 7.5, 150 mM NaCl, 0.5 mM EDTA, 0.5 % NP-40) supplemented with a protease inhibitor cocktail (Pierce Protease Inhibitor Tablets, ThermoFisher Scientific) for 30 min, spun at 15,000 × g for 10 min to clear the cell debris, and the cleared lysate was diluted with wash buffer (10 mM Tris/Cl pH 7.5, 150 mM NaCl, 0.5 mM EDTA). The resulting lysate was incubated with either GFP-Trap or anti-HA magnetic beads for 1 hour at 4°C. After incubation, the lysate was removed and the beads were washed with either wash buffer (for GFP-Trap beads) or TBST (TBS + 0.1% Triton X-100 for anti-HA beads). Bound proteins were eluted by boiling at 99°C for 10 min in 2× Laemmli buffer. The lysates (input) and eluted proteins were separated on a 12% SDS-PAGE gel (Bio-Rad) and transferred to a PVDF membrane for immunoblot analysis. The blot was probed with antibodies against HA (1:1000) and GFP (1:10,000).

For immunoprecipitation of mCherry-Lpg2411, HEK293T cells were transfected with plasmids encoding mCherry-Lpg2411 or mCherry alone using Lipofectamine 3000 reagent (ThermoFisher Scientific) according to the manufacturer’s instructions. mCherry-fusion proteins were isolated by immunoprecipitation using the RFP-Trap®_MA kit (Chromotek). Lysates from transfected cells were prepared as above and incubated with RFP-trap magnetic agarose beads for 30 min to capture mCherry or mCherry-Lpg2411. The beads were first washed in the wash buffer provided with the kit supplemented with a protease inhibitor cocktail (Pierce Protease Inhibitor Tablets, ThermoFisher Scientific). Finally, the beads were boiled in 2× Laemmli buffer to elute the proteins bound to the beads. The mCherry-fusion lysate (input) and proteins eluted from the beads were separated on a 4–20% TGX SDS-PAGE gel (Bio-Rad) and transferred to a PVDF membrane for immunoblot analysis. The immunoblot was probed with antibodies against ubiquitin (1:1000) and mCherry (1:1000).

### Assay for HA-Lpg2411 translocation timing

To determine timing of translocation of HA-Lpg2411 during infection, U937 macrophages were infected with a Δ*lpg2411* strain carrying a plasmid encoding HA-Lpg2411 at a multiplicity of infection (MOI) of 20. At the specified time points, cells were fixed as mentioned above, permeabilized with 0.1% Triton X-100 for 20 min and immunostained with rat anti-HA (1:1000) and Texas Red-conjugated goat anti-rat secondary antibody (1:3000). Bacteria were immunolabeled with anti-*Legionella* rabbit antibodies (1:6000) and an Alexa Fluor 488-conjugated anti-rabbit secondary antibody (1:3000). The percentage of infected cells harboring HA-signal was determined by scoring 100 cells per coverslip with three replicates for each time point. Microscopy was carried out using an epifluorescence microscope (ZEISS AxioObserver D1). This assay was repeated in three independent experiments.

### Infection of HT1080 cells for microscopy

HT1080 cells were grown in an 8-chamber cover glass (Cellvis #C8-1-N) pre-coated with collagen (Thermo Fisher #A1064401) according to the manufacturer protocol. Cells were challenged with *L. pneumophila* strains Lp02 (wild-type) or Lp03 (T4SS-defective mutant), thymidine-auxotroph derivatives of *L. pneumophila* strain Philadelphia-1 (Berger and Isberg, 1993). Bacteria were diluted in cell culture media and added to HT1080 cells at the indicated MOI, centrifuged for 5 min at 200 × g, and then incubated at 37°C with 5% CO_2_. If needed, after 2 hours of infection, the monolayer was washed three times with PBS to remove unbound bacteria. Cells were imaged with a laser scanning confocal Zeiss LSM 800 with Airyscan microscope with a 63× Plan-Apochromat 1.4 NA objective with Definite Focus. Live-cell imaging of bacterial uptake and replication in HT1080 cells was conducted on a laser-scanning confocal LSM880 microscope with a 63× Plan-Apochromat 1.4 NA objective with Definite Focus. Cells were imaged on a 37°C heated stage in normal growth media with 5% CO_2_ maintained by a CO_2_ chamber.

When needed, HT1080 cells were transiently transfected using Amaxa electroporation system according to the manufacturer protocol. Briefly, HT1080 cells were transfected using Amaxa Nucleofector Kit T (Lonza #VCA1003). For each transfection, 1×10^6^ cells were pelleted and resuspended in 100 µl room temperature Nucleofector Solution T. Two micrograms of total plasmid DNA was added, and cells were electroporated using Amaxa Program L-005 for high efficiency. Cells were transferred to collagen-treated coverglass and incubated for 4-6 hours before electroporation media was removed and cells were overlaid with fresh culture media. Cells plated on to collagen-treated coverglass were imaged 16-24 hours after electroporation. pCDNA3.1-YFP-KDEL was a kind gift from Dr. Jennifer Lippincott-Schwartz (Snapp et al., 2006). pEGFP-N1-LAMP1 was a kind gift from Dr. Juan Bonifacino (Farías et al., 2017). pEGFP-C1-Rab1a was a kind gift from Dr. Mitsunori Fukuda (Tohoku University) (Matsui et al., 2011). mEos-Calnexin was a kind gift from Dr. Michael Davidson (Addgene #57489). Ubiquitin staining was performed by fixing cells with 4% paraformaldehyde for 10 min, washing with PBS, permeabilizing with Triton X-100, then incubating with mouse anti-Ubiquitin (Sigma ST1200; 1:1000, 1 hour, 37°C), washing 3 times with PBS, then incubating with goat-anti-mouse-488 (Jackson Immuno Research #115-095-003; 1:1000, 1 hour, 37°C)

### Colocalization of HA-2411 and autophagic markers during host infection

To determine the localization of translocated HA-2411, HT1080 cells were infected with Δ*lpg2411* + pMMB207c-HA×4*-lpg2411* at an MOI of 40 for 15 hours. Cells were fixed and permeabilized with ice cold methanol or by incorporating 0.1% saponin in all staining steps. Cells were incubated with anti-HA (1:750) and either anti-LC3 (1:200), anti-p62 (1:100), or anti-FK2 (1:100) for 1 hour followed by either Texas Red– conjugated goat anti-rat (1:3000), Alexa Fluor 488–conjugated goat anti-mouse or anti-rabbit (1:3000). The nuclei and intracellular bacteria were stained with Hoechst (1:10,000) for 10 min. Coverslips were mounted with ProLong glass anti-fade mountant (ThermoFisher Scientific) and imaged using a confocal laser-scanning microscope (LSM880, Zeiss).

### Intracellular growth assay

U937 and HT1080 cells were seeded at a density of 2.5 × 10^5^ or 3 × 10^4^ cells/well in 24-well plates and infected with wild-type and mutant strains at an MOI of 0.05. Two hours after infection, wells were rinsed three times to eliminate extracellular bacteria. At 0, 48 and 72 hours post-infection, cells were lysed with 0.05% digitonin, diluted and plated on CYE plates. The number of bacteria recovered was recorded as colony forming units (CFU)/mL and standard deviations were calculated based on CFU values obtained from assays carried out in triplicate and over three independent experiments.

### Autophagic flux assay

Autophagic flux assay was performed as previously described (Choy et al., 2012). Briefly, HT1080 cells were infected with Lp01, Δ*lpg2411* + pMMB207c-HA×4 (empty vector), or Δ*lpg2411* + pMMB207c-HA×4*-lpg2411* at an MOI of 40 for 15 hours or left uninfected. Cells were then treated with 160 nM bafilomycin in DMEM (Sigma-Aldrich) or left untreated for 2 hours. Lysate was prepared using lysis buffer as described above. After boiling in Laemmli buffer, lysates were separated on a 12% SDS-PAGE gel (Bio-Rad) and transferred to a PVDF membrane for probing with anti-LC3 (1:1000), anti-ICDH (1:10,000), and anti-actin (1:5000). Western blot quantifications are representative of three independent experiments. ImageJ (version 1.53g) was used to quantify the band signal intensity on the western blot and a One-way ANOVA with Tukey’s multiple comparisons test was applied. Statistical data were calculated and graphed using GraphPad Prism5 (GraphPad, Inc., La Jolla, CA, USA).

## Supplemental Material

Transmission electron microscopy showed GFP-Lpg2411 is present on vesicular and elongated structures in CHO cells. A Lpg2411-GST fusion protein was generated to investigate direct lipid interactions between Lpg2411 and PIPs in a protein-lipid overlay assay. HT1080 human fibrosarcoma cells can be used as a model for studying *L. pneumophila* infection. A *L. pneumophila* strain harboring an IPTG inducible HA×4-Lpg2411 plasmid was generated to study the localization of Lpg2411 in host cells during infection. A *L. pneumophila* strain lacking *lpg2411* does not exhibit an intracellular growth defect in U937 and HT1080 cells.

## Abbreviations

HA: hemagglutinin

CFU: colony-forming units

CHO: Chinese hamster ovary cells

LCV: *Legionella*-containing vacuole

MOC: Mander’s Overlap Coefficient

MOI: Multiplicity of infection

PIP: phosphatidylinositol phosphate

PI(3)P: phosphatidylinositol-3-phosphate

PI(4)P: phosphatidylinositol-4-phosphate

T4SS: type IV secretion system

## Acknowledgements

This work was supported by NIAID, National Institutes of Health grant 1R21AI142317-01 (to M.R.N.), NSF CAREER award IOS-1750742 (to M.R.N.), Delaware COBRE Program, NIGMS, National Institutes of Health Grant P20GM104316 (to M. R. N.), as well as the Intramural Program of NICHD (grant 1ZIAHD008893-10 for S.S.L. and 1ZIAHD001609-26 for C.D.W). The LSM880 confocal microscope at the University of Delaware was acquired with a shared instrumentation grant (S10 OD016361) and access was supported by the NIH-NIGMS (P20 GM103446). The content is solely the responsibility of the authors and does not necessarily represent the official views of the National Institutes of Health or the National Science Foundation.

## Author Contributions

This study was directed by C. M. Pike, R. R. Noll, and M. R. Neunuebel and conceived and designed by C. M. Pike, R. R. Noll, and M. R. Neunuebel. C. M. Pike and R. R. Noll performed experiments. C.D.W. and S.S.L. established and validated the HT1080 cell line as a model for *Legionella* pathogenesis.

---

**Supplemental Table S1.**
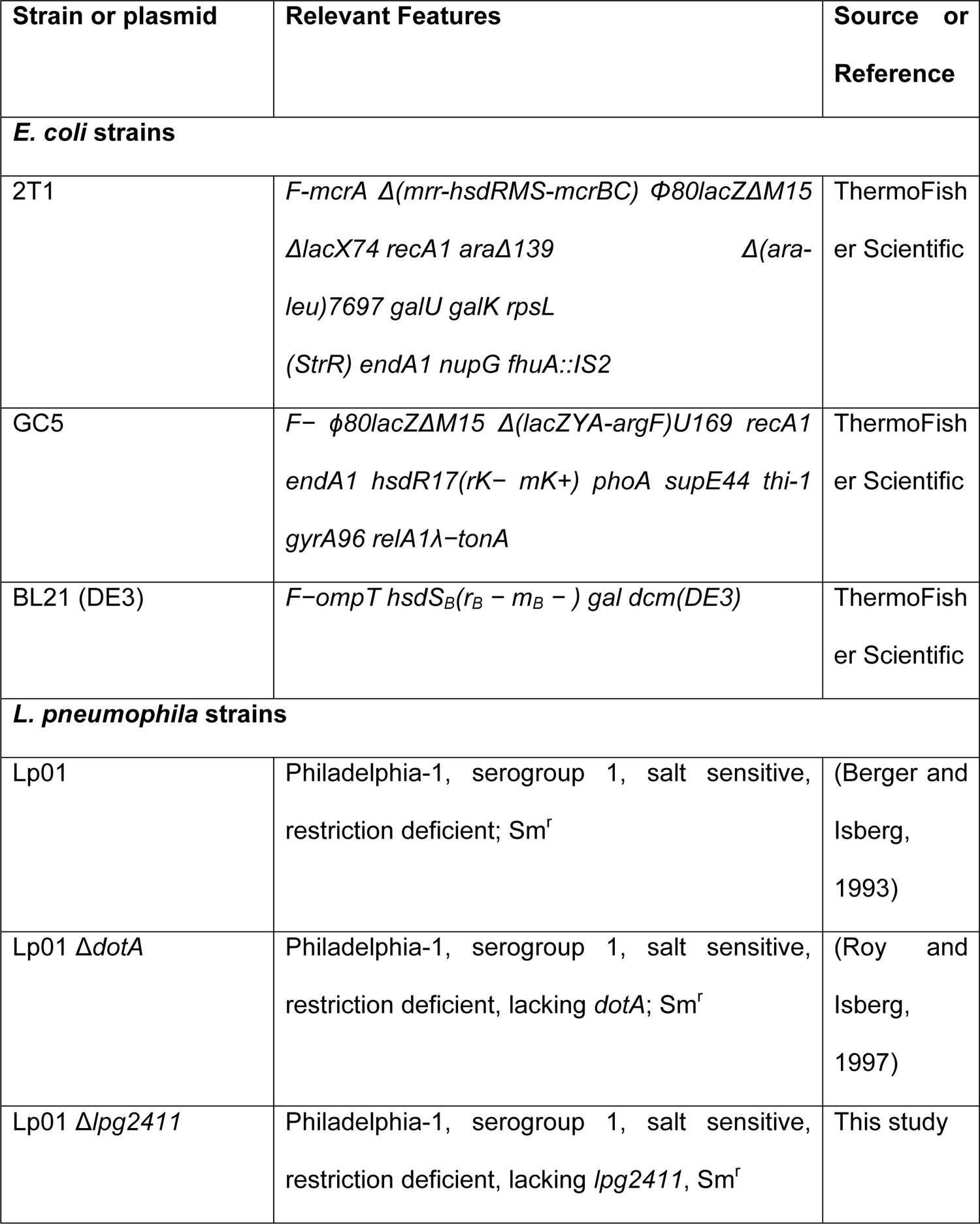
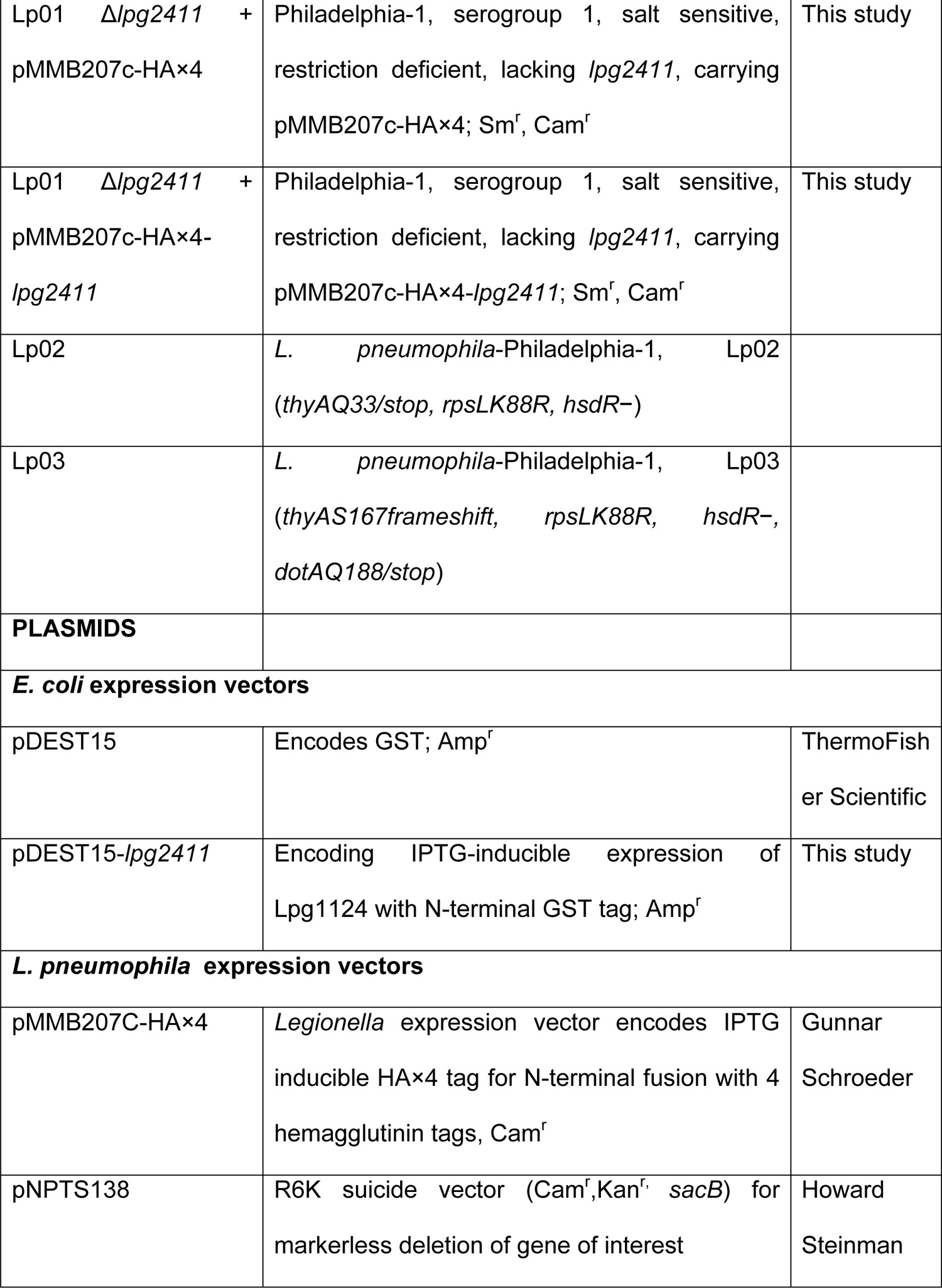
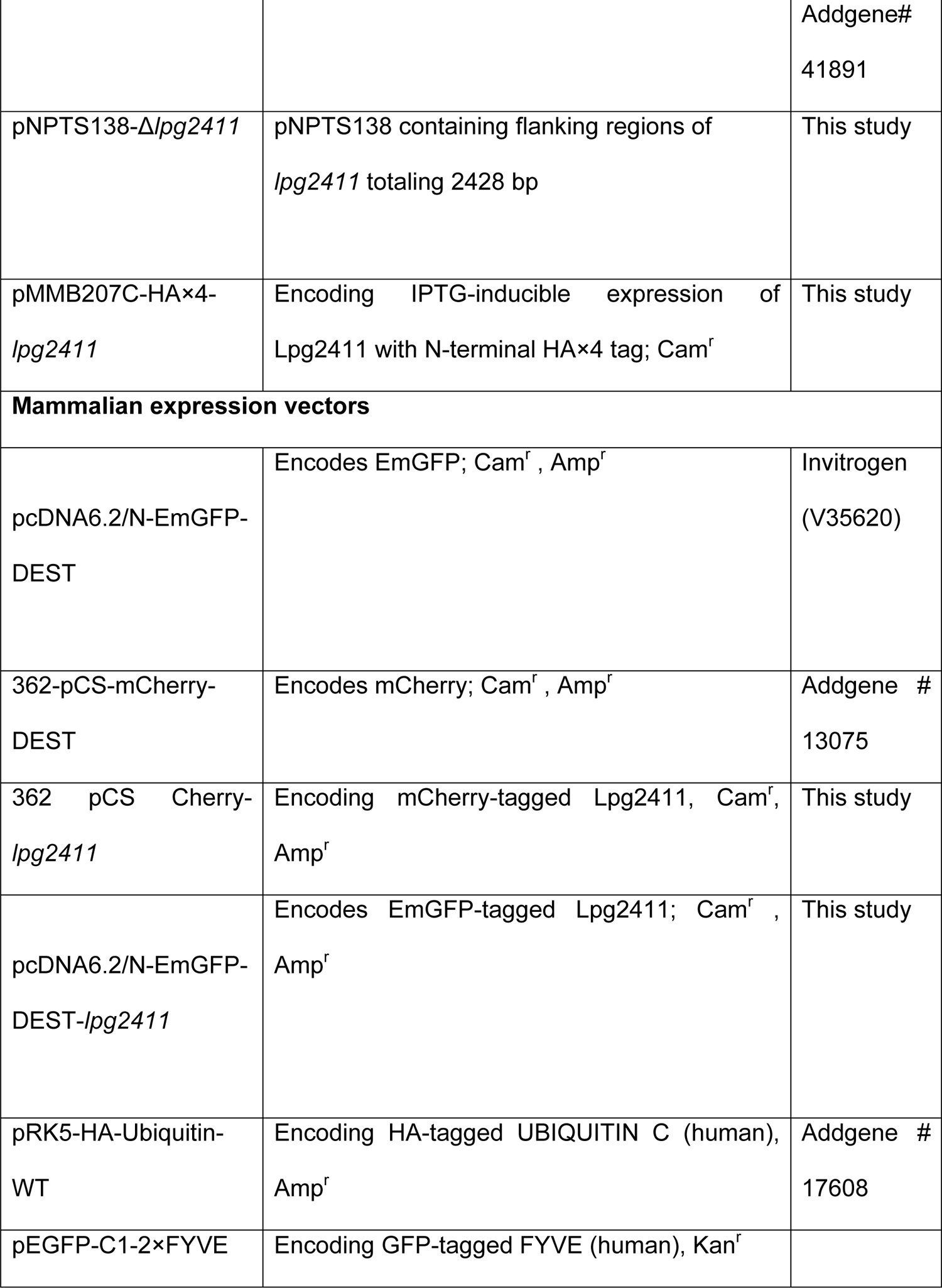
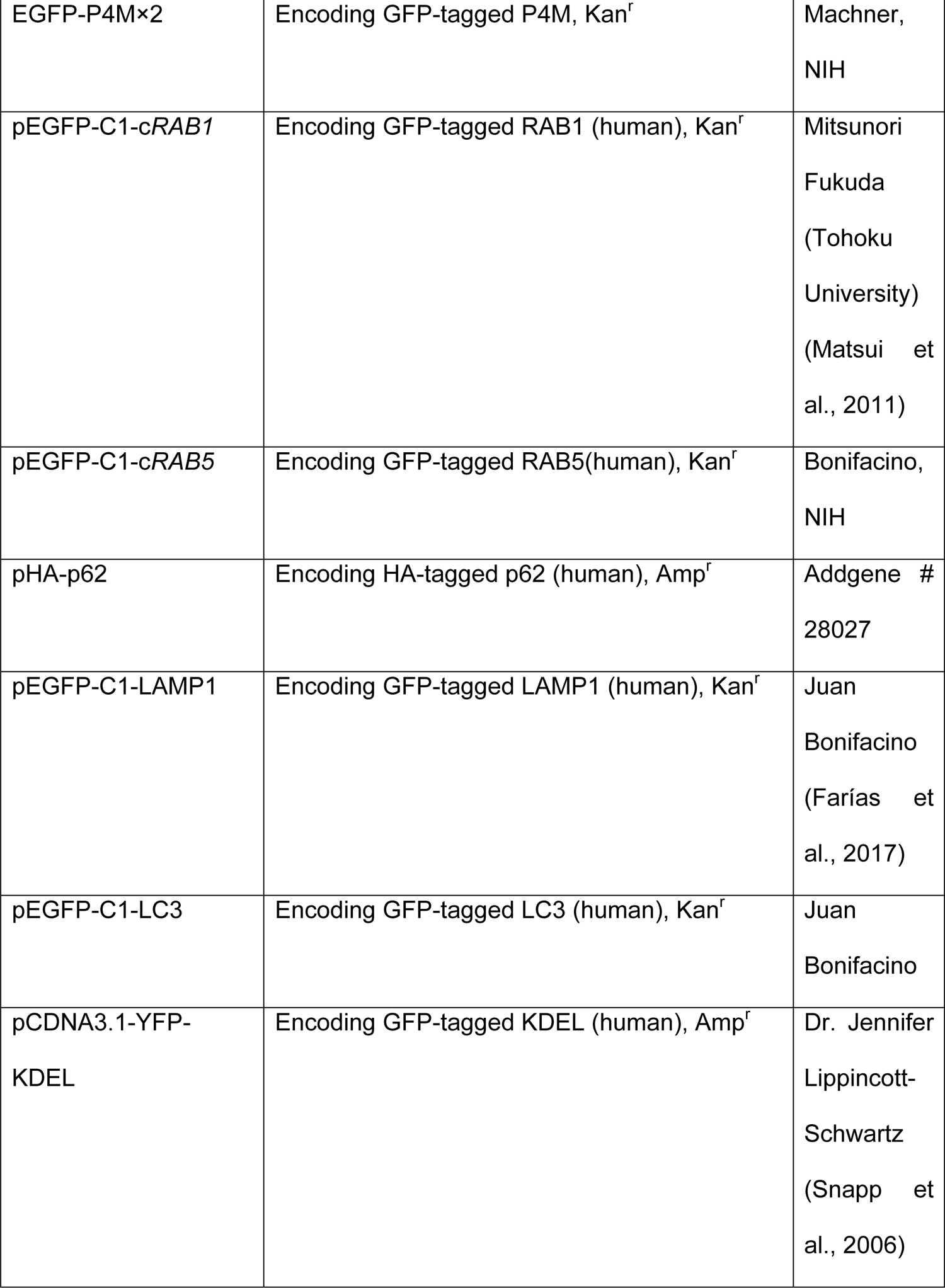
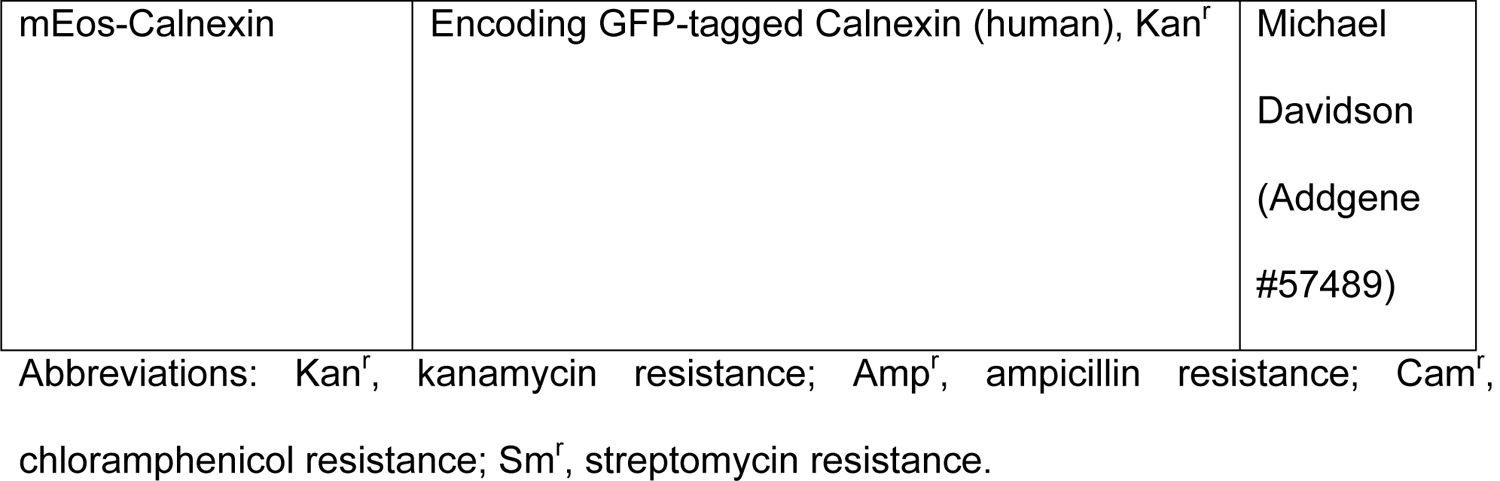
List of strains and plasmids used for this study.

**Supplemental Table S2.**
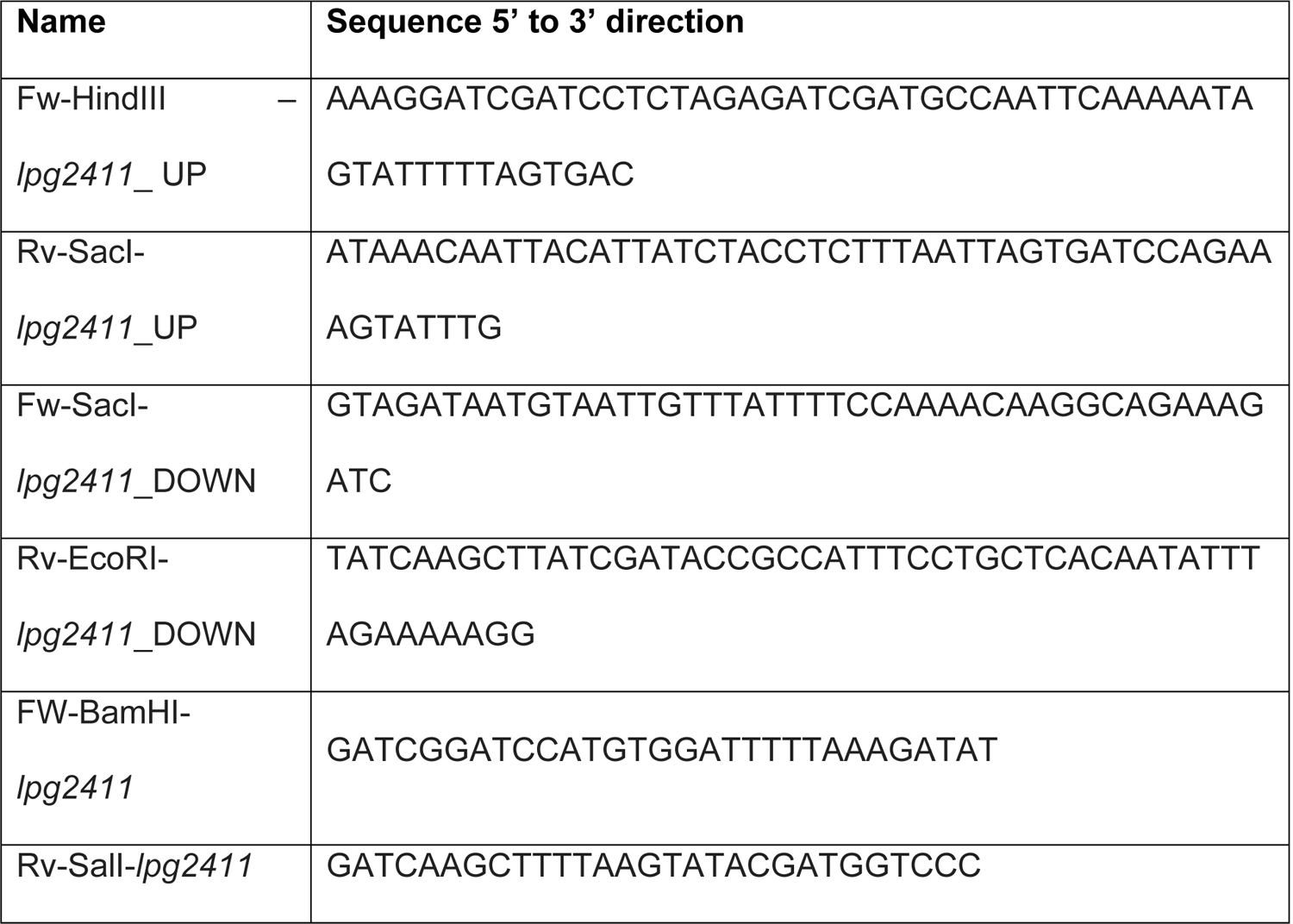
List of oligonucleotides used for this study.

**Supplemental Figure 1:**
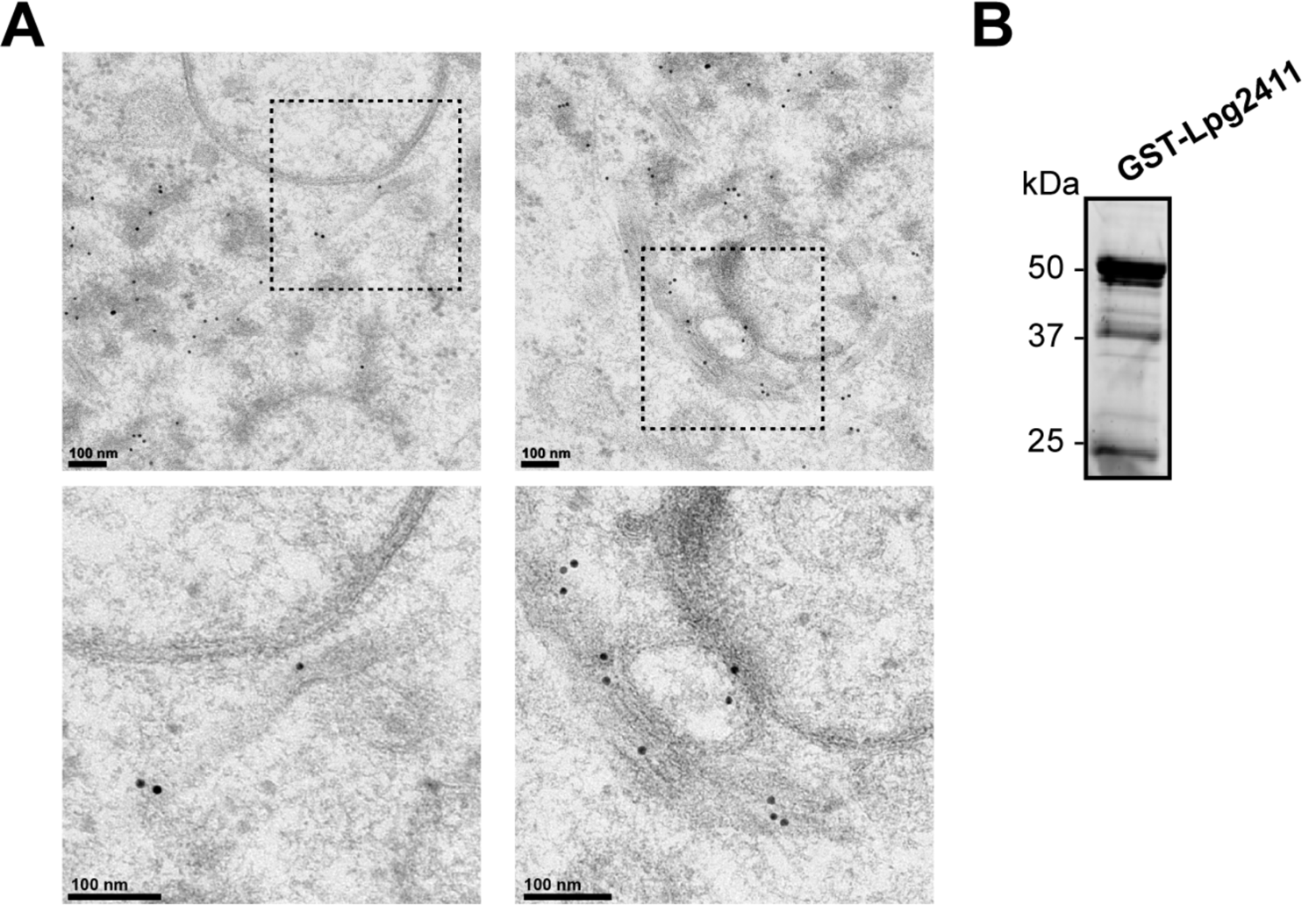
(A) Two representative transmission electron micrographs showing immunogold localization of GFP-Lpg2411 in CHO cells. Fixed cells were stained with a polyclonal rabbit anti-GFP antibody. Areas highlighted by rectangles (dashed line) in the top panels are magnified in the bottom panels. Scale bars, 100 nm. (B) Purified GST-Lpg411 (49 kDa) separated by SDS-PAGE on a 12% TGX stain-free gel.

**Supplemental Figure 2:**
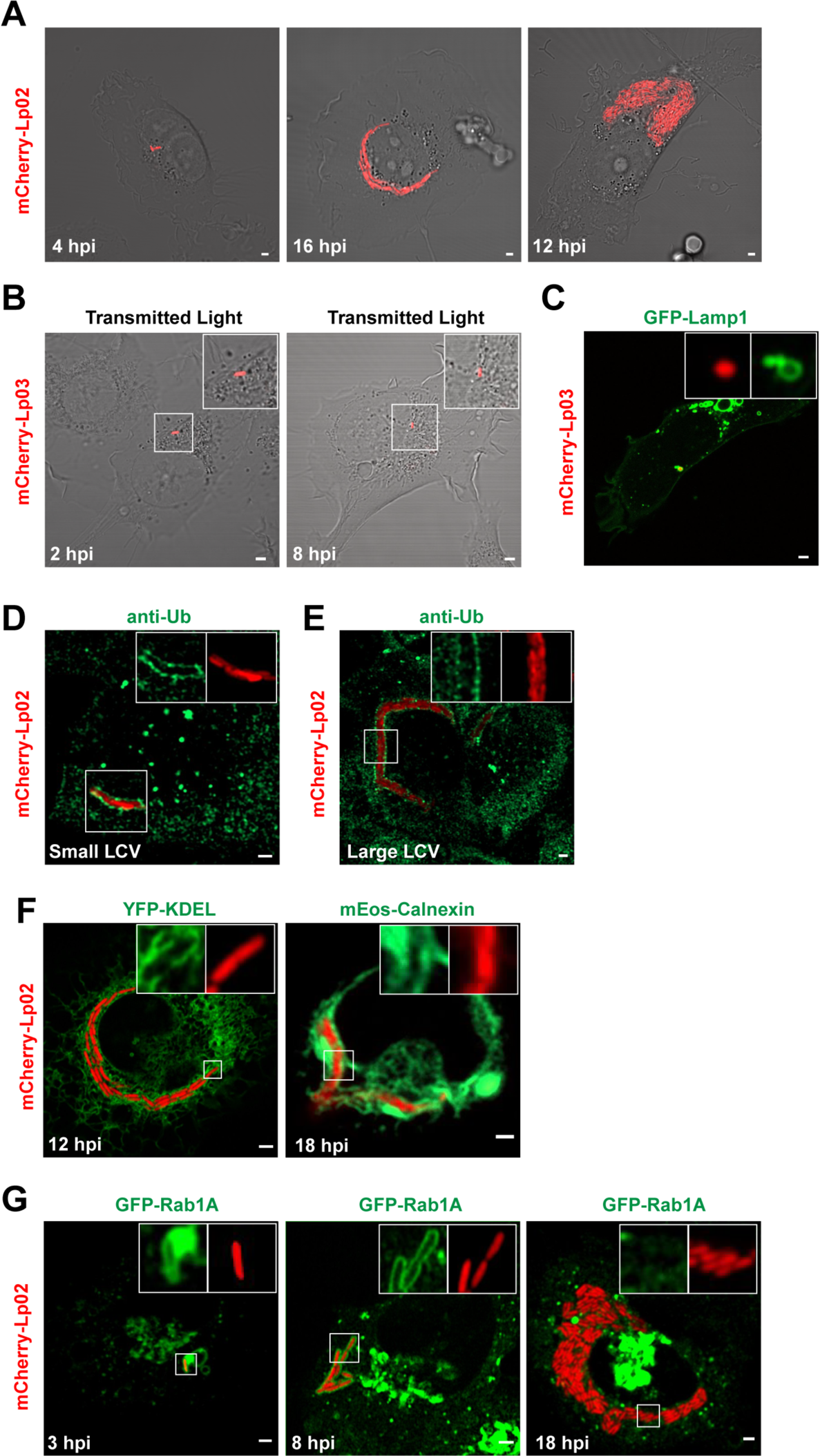
HT1080 cells were challenged with Lp02-mCherry or Lp03-mCherry as indicated, at MOI 10-25, for the indicated time points. (A) *L. pneumophila* infects and replicates in HT1080 cells. (B) Secretion system-defective Lp03 mutants enter HT1080 cells but do not replicate. (C) *L. pneumophila* Lp03 bacteria reside in GFP-tagged LAMP1 positive compartments. (D-E) *L. pneumophila* Lp02-containing vacuoles are decorated with ubiquitin. Cells were fixed and stained for ubiquitin and images show LCVs of various sizes. (F) LCVs co-localize with the ectopically expressed fluorescently-tagged ER markers YFP-KDEL and mEos-calnexin. (G) Rab1 is recruited to the LCV at early time points of infection and is removed from the LCV at later time points in HT1080 cells. Scale bar, 2 mm.

**Supplemental Figure 3:**
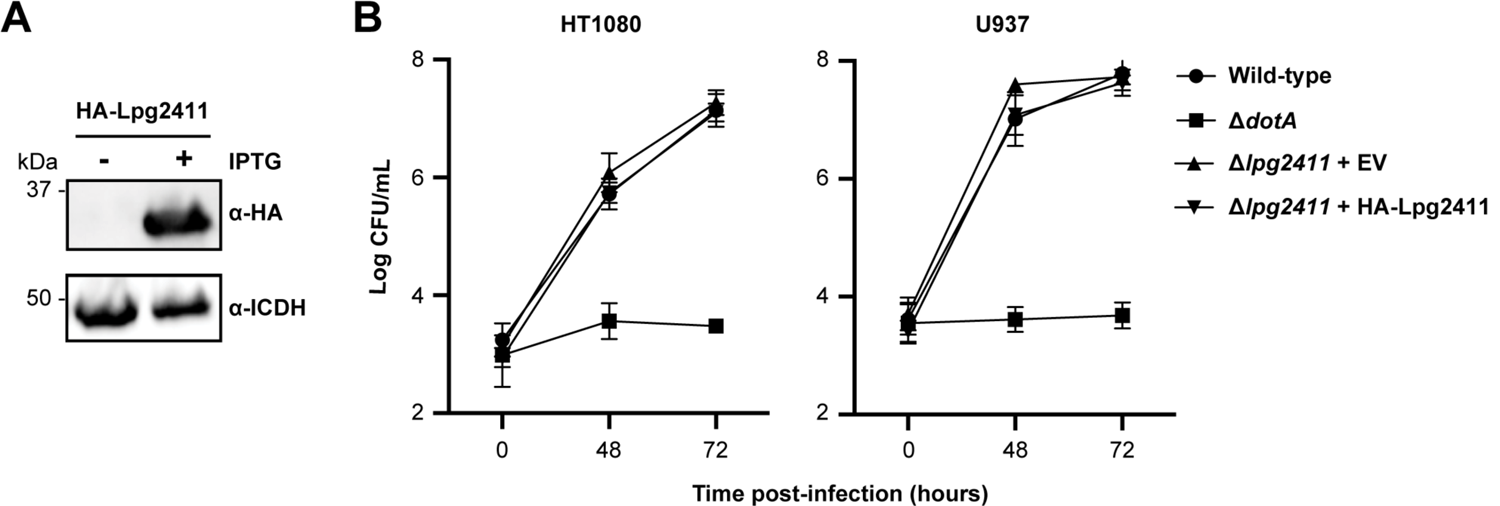
(A) Production of HA×4-Lpg2411 by the Δ*lpg2411* + pMMB207c-HA×4-*lpg2411* strain. *L. pneumophila* strain Δ*lpg2411* complemented with pMMB207c-HA×4-*lpg2411* was grown in the presence or absence of IPTG. Whole cell lysate was separated by SDS-PAGE electrophoresis on a 12% stain-free TGX gel and HA×4-Lpg2411 was detected by western blot with an HA antibody. (B) Lpg2411 is dispensable for intracellular growth. Monolayers of HT1080 cells or differentiated U937 monocytes were infected with wild-type, Δ*dotA*, Δ*lpg2411* + pMMB207c-HA×4, or Δ*lpg2411* + pMMB207c-HA×4-*lpg2411* at an MOI of 0.05, and cells were maintained at 37°C. The number of intracellular bacteria was determined by recording the number of colony-forming units per milliliter. The mean and standard deviation from three independent experiments is displayed for each time point.

**Supplemental Movie 1:** Membrane ruffling and uptake of GFP-Lp02 infected HT1080 cells. Fluid filled macropinosomes are easily identified in the transmitted light channel as large spherical vesicles within the cell which shrink over time. Internalized bacteria are not normally observed trapped within the lumen of macropinosomes, consistent with entry via phagocytosis. HT1080 cells were infected with Lp02-GFP, MOI 25. Imaging was started 0.5 hpi with images taken every 10 seconds for 11 min 40 seconds. Scale bar, 5 mm.

